# Emergent community agglomeration from data set geometry

**DOI:** 10.1101/109587

**Authors:** Chenchao Zhao, Jun S. Song

## Abstract

In the statistical learning language, samples are snapshots of random vectors drawn from some unknown distribution. Such vectors usually reside in a high-dimensional Euclidean space, and thus, the “curse of dimensionality” often undermines the power of learning methods, including community detection and clustering algorithms, that rely on Euclidean geometry. This paper presents the idea of effective dissimilarity transformation (EDT) on empirical dissimilarity hyperspheres and studies its effects using synthetic and gene expression data sets. Iterating the EDT turns a static data distribution into a dynamical process purely driven by the empirical data set geometry and adaptively ameliorates the curse of dimensionality, partly through changing the topology of a Euclidean feature space ℝ^*n*^ into a compact hypersphere *S*^*n*^. The EDT often improves the performance of hierarchical clustering via the automatic grouping information emerging from global interactions of data points. The EDT is not restricted to hierarchical clustering, and other learning methods based on pairwise dissimilarity should also benefit from the many desirable properties of EDT.

PACS numbers: 89.20.Ff, 87.85.mg

## I. INTRODUCTION

Community detection, better known as clustering in the literature of statistical learning [1–7], is a process of merging similar nodes of a complex network into communities (clusters) and often shows a hierarchical organization of communities at different levels of similarity. Akin to the idea of renormalization group in physics, decreasing the threshold for similarity leads to increasingly coarse-grained pictures of the “microscopic” network. The reduction in complexity can sometimes yield more interpretable statistical models that could serve as a basis for further classification analysis. Along this line, we present an idea of transforming dissimilarity measures to allow dynamic agglomeration of data points into communities.

Complexity in networks is analogous to that in many-body systems. Thus, clustering algorithms based on classical spin models have been designed by statistical physicists; e.g. each data point is replaced by a spin, and the similarity between points is computed from their Euclidean distance and spin orientations [8, 9]. Although such algorithms are both applicable and theoretically interesting, they usually require intensive Monte Carlo simulations and are thus too complex to implement in practical data analysis compared to other popular deterministic clustering algorithms. With practicality in mind, we present a nonlinear transformation of data set geometry and then pass the transformed geometric information to the standard hierarchical clustering algorithm widely used in contemporary data analysis. We show that the geometric transformation effectively captures a collective interaction among all sample points and that such global network interaction often improves the accuracy of community detection.

Most statistical learning algorithms utilize a pairwise dissimilarity measure 
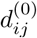
 that depends only on the (*i*, *j*)-pair of samples. In the context of Einstein’s theory of gravity, or Riemannian geometry, the “geometry” is completely encoded in the metric tensor. This paper adopts the same notion of geometry for a data set and focuses on the information encoded in the dissimilarities between all pairs of sample points. The *n* features of *m* samples measured in an experiment are typically organized into an *n* × *m* matrix, with *m* samples represented as points in ℝ^*n*^. Thus, the Euclidean *L^p^*-metric directly defined on the feature space ℝ^*n*^ is among the most common pairwise dissimilarities.

In high dimensions, however, the relative contrast between the farthest and nearest points measured by the *L^p^*-metric diminishes; consequently, the concept of nearest neighbors, which serves as the foundation for clustering, becomes increasingly ill-defined as the feature dimension increases [10–12]. This phenomenon is termed “the curse of dimensionality,” analogous to the idea of “more is different” for many-body systems [13]. Modifications of Euclidean distances are found to improve the relative contrast for an artificial data cloud drawn from a single distribution [10, 11], but fail in data drawn from several distributions [12]. One way to address the loss of contrast in high dimensions for multi-distribution data is to introduce an effective dissimilarity measure calculated from the number of shared nearest neighbors of two data points, where each point is allowed to have a fixed number of nearest neighbors [12]. The use of effective dissimilarity reduces the effect of high feature dimensions in subsequent computations; however, the choice of effective dissimilarity function actually dictates the improvement.

This paper proposes a new effective dissimilarity transformation (EDT), where all data points in the primary feature space participate in redefining the effective dissimilarity between any two given data points. Our main motivation stems from the empirical formula for distance correlation in statistics and the idea of heat flow on a hypersphere in support vector machine classification [15, 16]. Empirical distance correlation utilizes the co-variance of pairwise distance between all samples to measure statistical association between two random vectors [14]. In this spirit, our EDT can be viewed as measuring the similarity between two data points by taking a dot product of the corresponding columns of uncentered distance matrix. More precisely, transforming data to lie on a hypersphere has been previously shown to yield several advantages in machine learning [15, 16], so an intermediate step in EDT maps the columns of distance matrix to points on a hypersphere before taking a dot product. We show that this simple EDT improves the contrast between clusters in a geometrically interpretable manner and that it is able to reshape a geometrically mixed data distribution into separable clusters.

To be specific, the effective dissimilarity obtained from EDT is beyond the pairwise level and globally captures relations to all available sample points. Moreover, we also observe that the EDT is a map defined on a non-negative dissimilarity space 
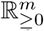
 of samples, where typically the sample size *m* is much smaller than the feature dimension *n*, thus providing an efficient dimensional reduction scheme. Iteratively applying the transformation yields a sequence 
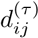
 of EDT parametrized by a non-negative integer *τ*. As *τ* increases, microscopic structures condense locally, while inter-cluster macroscopic distinctions become more evident. Since the heat kernels describing heat diffused from a point source is parametrized by continuous time *t* ≥ 0, we may interpret EDT as a generalized nonlinear diffusion process in the dissimilarity space driven by the distribution of samples. Iterating EDT thus turns a static distribution of points into a dynamical process and often amplifies its power of cluster separation.

## II. RESULTS

### A. Formulation of effective dissimilarity transformation (EDT)

As observed in previous support vector machine (SVM) classification studies [15, 16], hyperspherical geometry often improves classification accuracy. Motivated by these results, we now introduce an effective dissimilarity transformation based on a hyperspherical representation of data clouds. To map sample points onto a hypersphere, we will utilize the following hyperspherical transformation from non-negative space 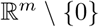 to a unit hypersphere:

#### Definition 1

*A hyperspherical projective map* 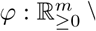 {0} → *S*^*m*−1^ *maps a vector* x, *with x_i_* ≥ 0 *and* 
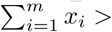
, *to a unit vector* 
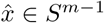
 *where* 
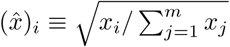
.

A useful measure of similarity on a hypersphere is the cosine similarity:

#### Definition 2

*For unit vectors* 
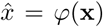
 *and* 
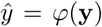
 *Obtained from non-negative vectors* 
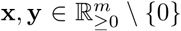
 *via the hyperspherical projective map, the cosine similarity is the dot product* 
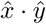
.

The EDT relies on this notion of cosine similarity, as explained below.

Many algorithms – such as hierarchical clustering, KMedoids, and KMeans – directly rely on some notion of difference between samples. For example, the Euclidean distance function is a popular measure of the difference between two sample points in ℝ^*n*^. In statistical learning approaches based on pairwise differences, however, we often relax the definiteness condition and triangular inequality satisfied by a distance function and utilize instead a more general and flexible measure of difference, called the dissimilarity function:

#### Definition 3

*A dissimilarity function defined on a manifold* 
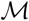
 *is a map d*: 
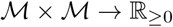
 *satisfying*

1. *non-negativity: d*(*x*, *y*) ≥ 0 *for all x*, 
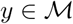
,
2. *identity: d*(*x*, *x*) = 0 *for all* 
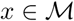
,
3. *symmetry: d*(*x*, *y*) = *d*(*y*, *x*) *for all x*, 
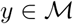
.

Usually 
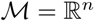
, representing the sample space of original data directly collected from experiments, and its nonlinear embedding into an abstract manifold is often only implicitly defined through the dissimilarity function.

Dissimilarity functions are relatively easy to construct; in particular, we can turn the cosine similarity on 
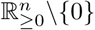
 into a dissimilarity function by defining 
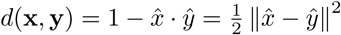
. We here show that this cosine dissimilarity function can be iteratively applied to an initial dissimilarity measure and that this simple iteration leads to several robust properties desirable for clustering applications.

More precisely, given an initial dissimilarity function *d*(·, ·) and *m* sample points, organize the pairwise dissimilarity of the samples into an *m* × *m* non-negative, symmetric dissimilarity matrix *d*^(0)^. To apply our method, we only need to assume the mild condition that each column of *d*^(0)^ is not a zero vector. We then define the effective dissimilarity transformation on the space of such matrices as follows:

**Figure 1.**
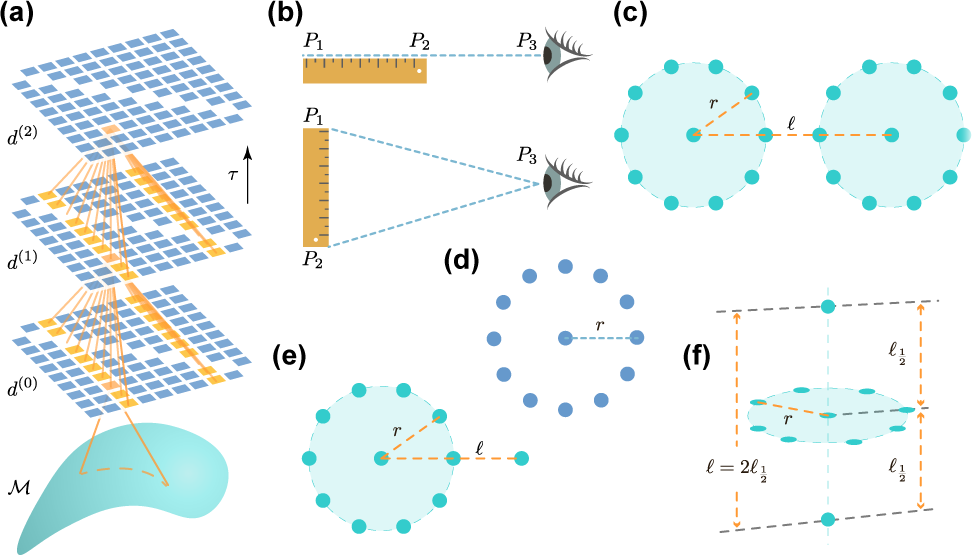
(a) A schematic illustration of the network structure of effective dissimilarity transformations (EDT) parameterized by *τ*. The (*i*, *j*)-th entry of *d*^(*τ*)^ arises from transforming the *i*- and *j*-th columns of *d*^(*τ* − 1)^. (b) Illustrations of *perspective contraction* effect of EDT. (c) Two ideal clusters with radius *r* and centroid-centroid distance *l* in ℝ^2^. (d) The detector used in the measurement of *local deformation* of data distributions in ℝ^2^. (e) An ideal cluster of radius *r* in ℝ^2^ and an outlier at distance *ℓ* from the cluster centroid. (f) An ideal cluster of radius *r* in the *xy*-plane of ℝ^3^ with symmetrically located outliers on the *z*-axis at distance 
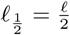
 from cluster centroid.

**Figure 2.**
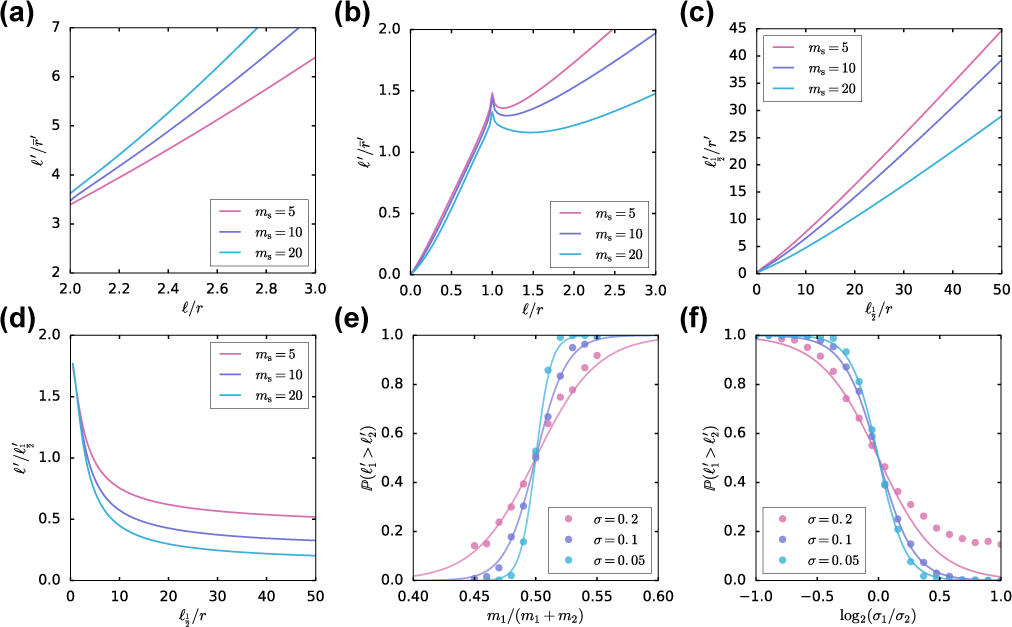
Results of Gedankenexperimente: (a) cluster condensation (Fig. 1(c)), (b) single outlier absorption (Fig. 1(e)), (c, d) two outliers perpendicular to an ideal cluster (Fig. 1(f)), (e, f) probabilistic sampling.

#### Definition 4

*The effective dissimilarity transformation (EDT)* 
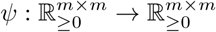
 *is defined as*

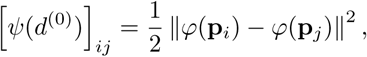

*where* **p**_*i*_ *is the i-th column of the dissimilarity matrix d*^(0)^ *and φ is the hyperspherical projective map into S*^*m*−1^. *We denote d*^(1)^ ≡ *ψ*(*d*^(0)^).

The resulting *d*^(1)^ is thus a cosine dissimilarity matrix of the *m* samples newly represented by the columns of the dissimilarity matrix *d*^(0)^. Importantly, the pairwise dissimilarity captured by *d*^(1)^ between any two samples measures how dissimilar are their respective *d*^(0)^ dissimilarities to all samples; in other words, each entry of *d*^(1)^ depends on the global network structure encoded in *d*^(0)^ as illustrated in Fig. 1(a). Iterating the map composition 
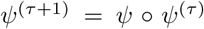
 yields a sequence 
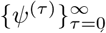
 of EDTs and corresponding dissimilarity matrices 
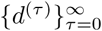
, where *ψ*^(0)^ is the identity map and *d*^(*τ*)^ = *ψ*^(*τ*)^(*d*^(0)^). The sequence of dissimilarity matrices 
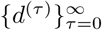
 may be interpreted as inducing a data-driven evolution or flow of sample points parametrized by *τ*. This paper shows that the data-driven redefinition of dissimilarity resulting from an iterated application of EDT often leads to improved clustering results.

Even though EDT is simple in its definition and deterministic in nature, its nonlinearity makes the flow of data points difficult to study. Consequently, we first study the properties of EDT by performing Gedankenexperimente on carefully designed synthetic data sets shown in Fig.1(b-f) (accompanying simulation results in Fig. 2a-f)), and then test the power of these observed properties in the setting of real data sets.

### B. Gedankenexperimente of EDT

First consider the simple data set consisting of 3 distinct points, *P*_1_; *P*_2_; and *P*_3_, in ℝ^*n*^, for any *n* ≥ 2. Let *P*_1_ and *P*_2_ represent two ends of a ruler of length 
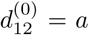
, and let *P*_3_ represent an observer at distance *b* to the center of the ruler; Fig. 1(b) shows two particular cases: (1) the ruler and observer are colinear, and *b* > a/2; (2) the observer and ruler form an isosceles triangle, and 
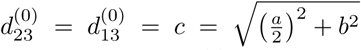
. In scenario (1), the original distance 
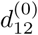
 between *P*_1_ and *P*_2_ is equal to the ruler length and is also the observed distance 
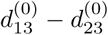
 measured by the observer at *P*_3_, irrespective of the location of *P*_3_; after EDT, however, both 
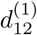
 and the ratio 
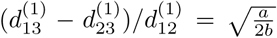
 shrink as the observer moves away (Appendix B1). That is, in the limit *b* ≫ *a*, the effective dissimilarity between *P*_1_ and *P*_2_ approaches zero, and the observer at *P*_3_ cannot distinguish between *P*_1_ and *P*_2_ on the scale set by 
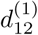
. In the language of hierarchical clustering, the single, average, and complete linkages become equivalent after EDT as *P*_3_ becomes a clear outgroup. Similarly, in scenario (2), the effective ruler length also shrinks as the observer moves away from the other two points, i.e. 
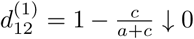
 as *b*/*a* ↑ ∞. We can thus summarize these properties as a *perspective contraction* effect:

**Observation 1** *The EDT dissimilarity between each pair of points shrinks as an observer moves away from the distribution of points. Consequently, compared to the original dissimilarity, hierarchical clustering using the EDT dissimilarity is insensitive to the choice of linkage*.

We verified this observation by comparing the performance of Euclidean distance with its EDT dissimilarity in the hierarchical clustering of three Gaussian clouds in ℝ^2^ using single, average and complete linkages (Fig. 3). As often is the case with real data, the three linkages based on the Euclidean distance led to different clustering results (Fig. 3 top row), whereas the EDT dissimilarity was insensitive to the choice of linkage (Fig. 3 bottom row).

**Figure 3.**
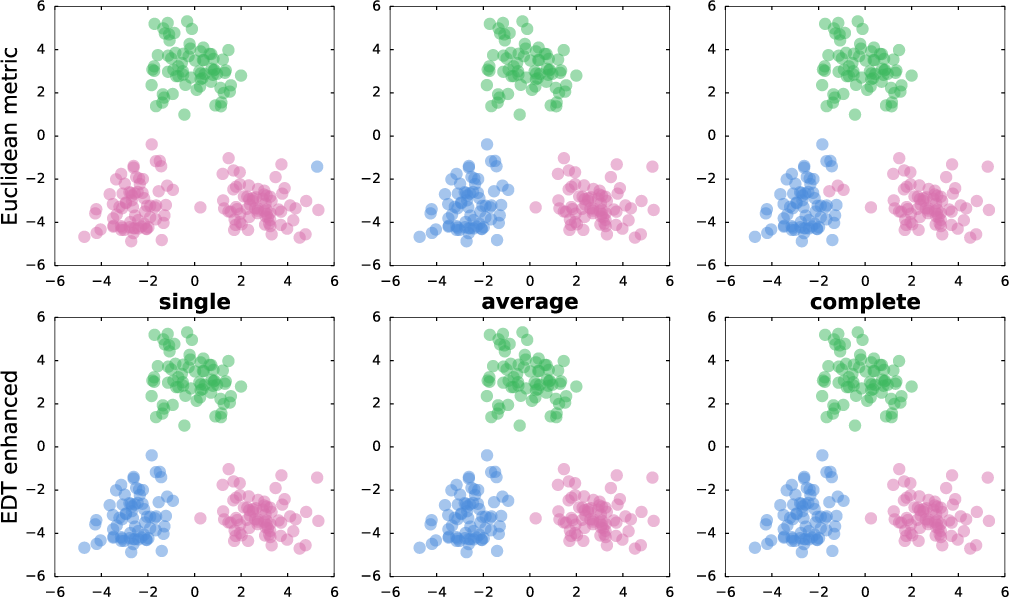
Comparison of hierarchical clustering results using Euclidean distance vs. EDT-enhanced Euclidean distance with single, average, and complete linkages. The number of clusters was chosen to be three in the analysis.

We next replaced the ruler and observer in our first model with two identical ideal clusters, each of which consisted of a centroid point and *m_s_* uniformly distributed satellites at radius 
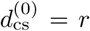
 in ℝ^2^(Fig.1(c)). The distance between the two centroids was set to 
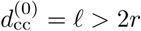
, and data distribution had two global mirror reflection symmetries about (1) the line connecting two centroids, and (2) the perpendicular bisector thereof. We compared the changes in intra- and inter-cluster dissimilarities after EDT and found that the two circles were deformed, but the global mirror reflection symmetries were preserved. We further measured the mean 
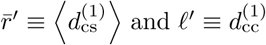
 and found the ratio 
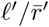
 to be an increasing function in both *ℓ*/*r* and *m_s_*; moreover, 
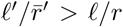
 for any *ℓ* > 2*r* (Fig.2(a)). Thus, the EDT had the effect of forcing the data points in each cluster to condense towards their respective centroid location, a potentially desirable effect that can help automatically merge data points into correct communities. We summarize our observation as a *cluster condensation* effect:

**Observation 2** *For separable clusters, the EDT condenses the points within a cluster, while inflating the space between clusters; this cluster condensation effect becomes stronger with the number of points in each cluster and also with the initial inter-cluster dissimilarity*.

The previous two Gedankenexperimente were performed on highly symmetric data sets. To probe the local deformation induced by EDT on a generic data distribution, we devised a detector, or a composite “test charge.” The idea is generalizable to higher feature dimensions, but to simplify the interpretation, we performed the simulation in ℝ^2^, with the detector being an ideal cluster of 12 sensor points at radius *r* from a centroid point (Fig. 1(d)). Deviations of the detector from a perfect circle in local ambient distributions were used to assess the EDT impact landscape. We captured the deviations through the transformed arm lengths 
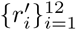
 of the 12 sensors after EDT; we then derived two scalar quantities of interest: (1) the mean arm length 
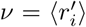
 that measures a volume change, and (2) the standard deviation of 
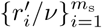
, denoted *κ*, that measures anisotropy or the effect of “tidal force” from probed data points. The observed volume changes were consistent with the effect of “perspective contraction,” and the mean arm length *ν* of the detector shrank as it moved away from high density regions of the probed data distribution (Fig. 4). The *κ*-distributions were highly non-trivial, as illustrated in Fig. 5: *κ* attained high values whenever the rim of the detector was near a data point, indicating an intense tug-of-war between the data points and the detector that were both trying to capture the sensors; by contrast, the normalized *κ* almost vanished at the centers of two Gaussian distributions, within the inner circle of the two layers of circularly distributed points, and at the center of “O” in the “COS” data. The low values of *κ* in the interior of Clustered data suggest a screening effect that shields the interior from anisotropic distortions, akin to the shielding effect of conductors in electrostatics; this effect may potentially protect sub-cluster structures within a dense cluster.

**Figure 4.**
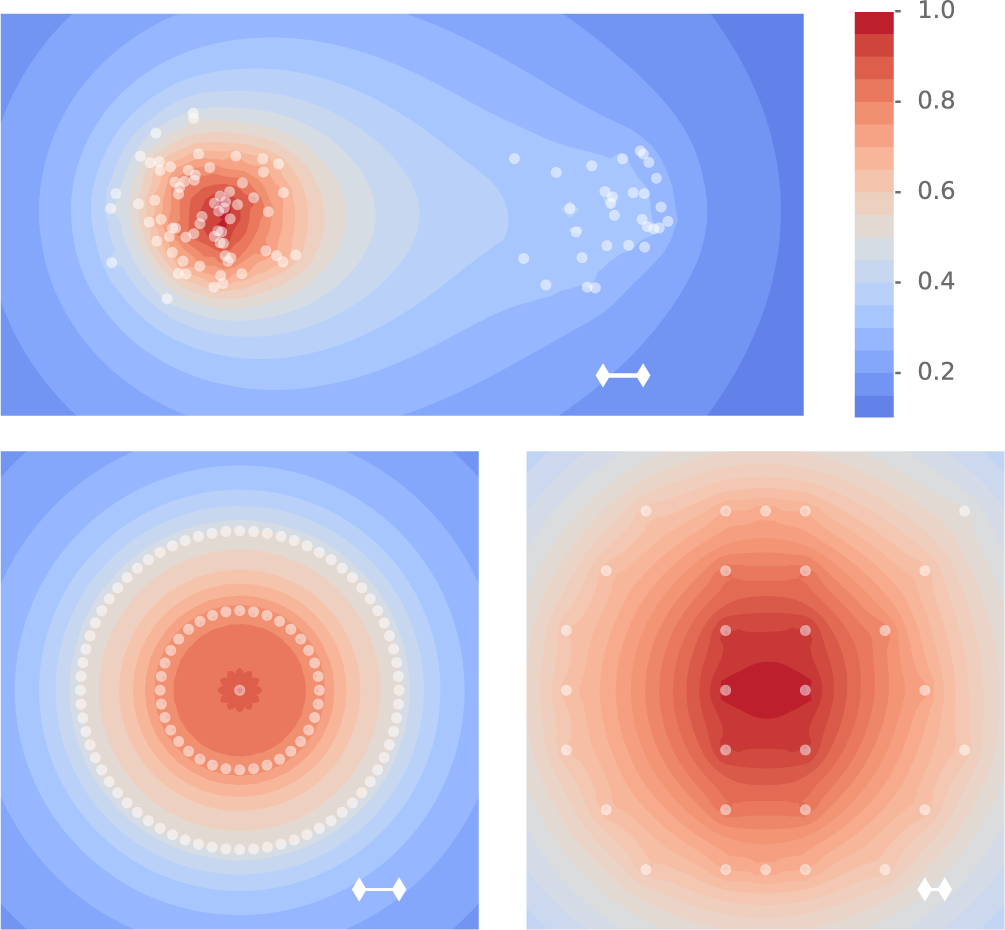
The ν-distribution for three data sets: (1) two Gaussian distributions with equal variance, but different sample sizes *m*_left_ = 70 and *m*_right_ = 30 (top); (2) two layers of circularly distributed points with radius *r*_outer_ = 2*r*_inner_ (bottom left); (3) points distributed in the shape of the word “COS” (bottom right). Each *κ*-distribution was normalized by dividing by its maximum; the white segment in each plot indicates the diameter of the detector used in the measurement of *ν*.

**Figure 5.**
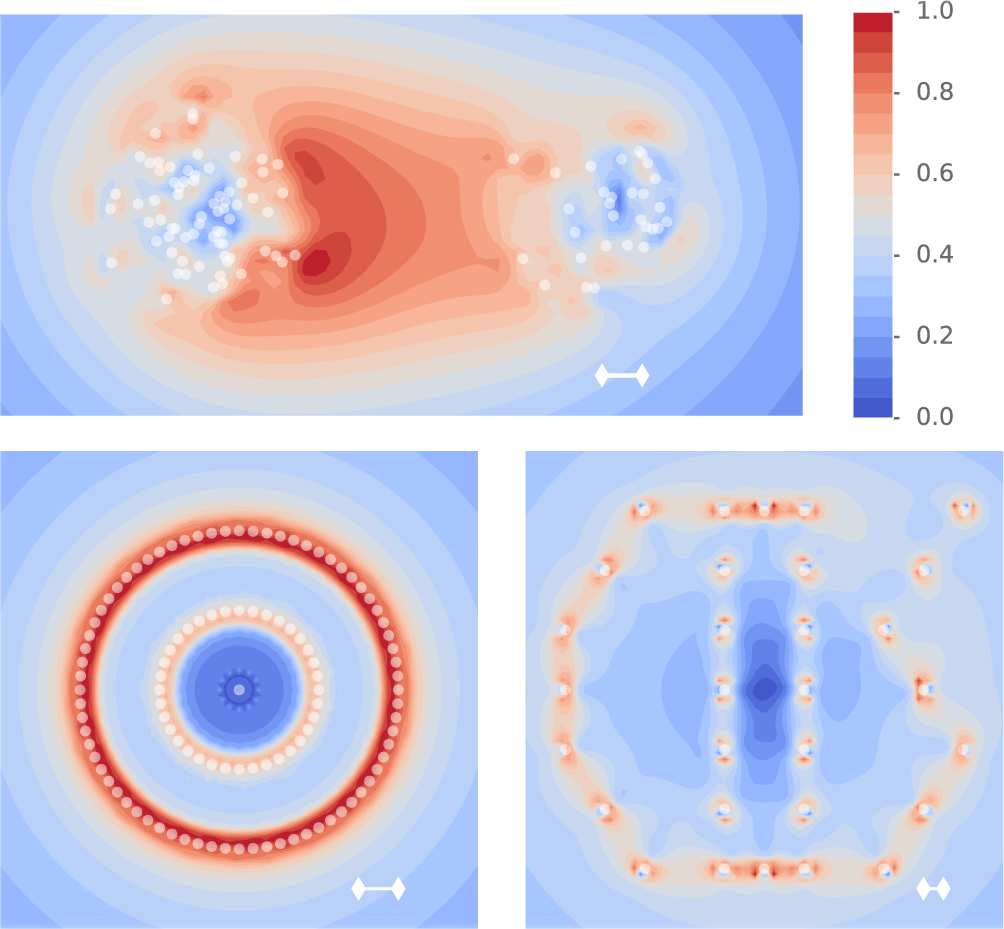
The *κ*-distribution for three data sets: (1) two Gaussian distributions with equal variance, but different sample sizes *m*_left_ = 70 and *m*_right_ = 30 (top); (2) two layers of circularly distributed points with radius *r*_outer_ = 2*r*_inner_ (bottom left); (3) points distributed in the shape of the word “COS” (bottom right). Each *κ*-distribution was normalized by dividing by its maximum; the white segment in each plot indicates the diameter of the detector used in the measurement of *κ*.

Inspired by the high values of *κ* near the boundary of a cluster, we performed additional experiments to test the effect of EDT on outliers, using (1) an ideal cluster in ℝ^3^ with *m_s_* satellites at radius *r* from the center point and an additional single point at varying distance *ℓ* from the center (Fig.1(e)), and (2) the same ideal cluster in the *xy*-plane of ℝ^3^ and two outliers located on the *z*-axis at *z* = ±*ℓ*/2 (Fig.1(f)). For the first case, Fig. 6 shows how a cluster of points traps an outlier and prevents it from escaping the cluster. Furthermore, in both cases, we observed that the trapping power increased with cluster mass *m_s_*: in case (1), increasing *m_s_* reduced the relative effective outlier-centroid dissimilarity 
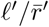
 and broadened the outlier region that got pulled back towards the cluster (Fig. 2(b)); in case (2), increasing *m_s_* also decreased the relative effective outlier-centroid dissimilarity 
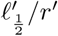
 Fig. 2(c)). We summarize the local deformation effect, or the “tidal force” exerted by local data distribution, as follows:

**Observation 3** *Under the EDT, data points deform the local distribution of neighboring points such that potential outliers tend to be trapped by a massive cluster. The deformation is strong near the exterior of a cluster and almost completely screened inside the cluster*.

**Figure 6.**
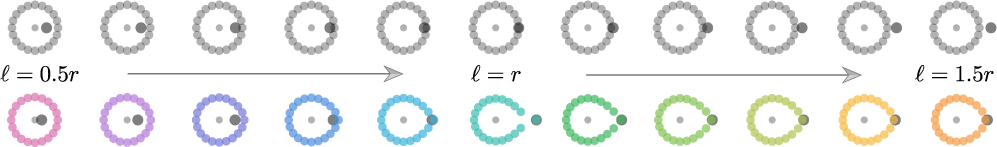
A cluster of points can pull back or “trap” an outlier. Figure shows the case illustrated in Fig.1(e) for varying values of the ratio *ℓ*/*r* in the range [0.5; 1.5] and for 20 satellite points. The top gray circles indicate the actual locations of points in ℝ^2^; the bottom colored circles illustrate the corresponding effective locations after EDT, where we doubled the distortions to visualize the effect more clearly. As *ℓ*/*r* increased from left to right, the deformed circle behaved like an elastic membrane trying to trap the outlier from escaping and demonstrated singular behavior at *ℓ* = *r*.

In case (2), we also observed an intriguing paradox: the transformed outlier-outlier dissimilarity *ℓ*' satisfied the condition 
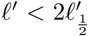
 for all 
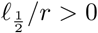
, and it even satisfied the counter-intuitive inequality 
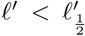
 for sufficiently large 
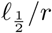
 and large *m_s_* (Fig. 2(d)). A resolution of this paradox is achieved by noting that the points at infinity become identi ed under EDT. For example, for the particular case of circularly distributed data points in ℝ^2^, as illustrated in Fig. 7, the outer rings of points become increasingly similar as *τ*, indexing the EDT iteration, increases; moreover, the effect becomes more pronounced as the density of points at the center of the distribution increases (bottom row in Fig. 7, Appendix B 4). In mathematical terms, adding the point at infinity to ℝ^2^ yields a compact sphere *S*^2^, and the above process can be visualized as the outer rings diffusing towards the south pole (Fig. 7).

**Figure 7.**
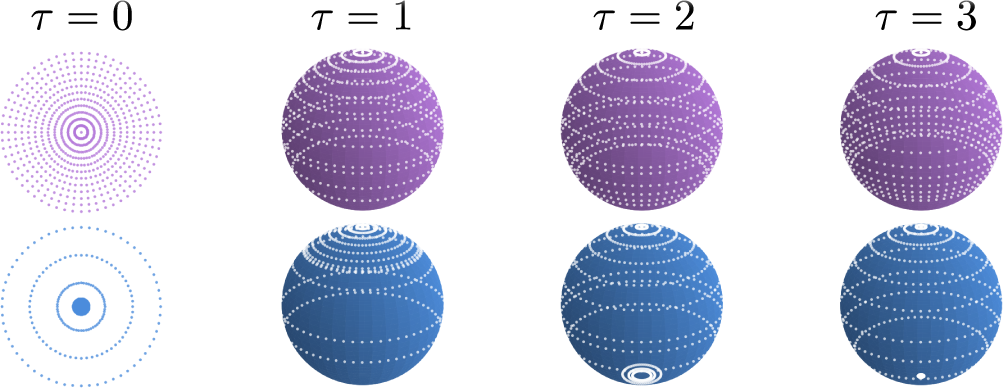
EDT approximately identifies the points at infinity. We designed two uniformly circularly distributed data sets with (1) a uniform increment in radius, or (2) a small increment in radius near the center and a large increment for the outermost three circles. For both data sets, the outer circles became relatively closer as *τ* increased. The effect was more pronounced in the second case, and the outermost three circles were visibly mapped to the south pole. The mapping method can be found in Appendix B 4.

We tested whether this property of EDT can help improve clustering performance on synthetic data sets that are known to confound simple algorithms. For this purpose, we chose two clusters of data concentrically distributed with a gap in radial direction (Fig. 8). The EDT dramatically improved the performance of hierarchical clustering with Euclidean metric (Fig. 8); furthermore, the EDT-enhanced hierarchical clustering outperformed spectral clustering using Gaussian RBF as a measure of similarity (Fig. 9). These observations can be summarized as EDT’s *global deformation* effect:

**Figure 8.**
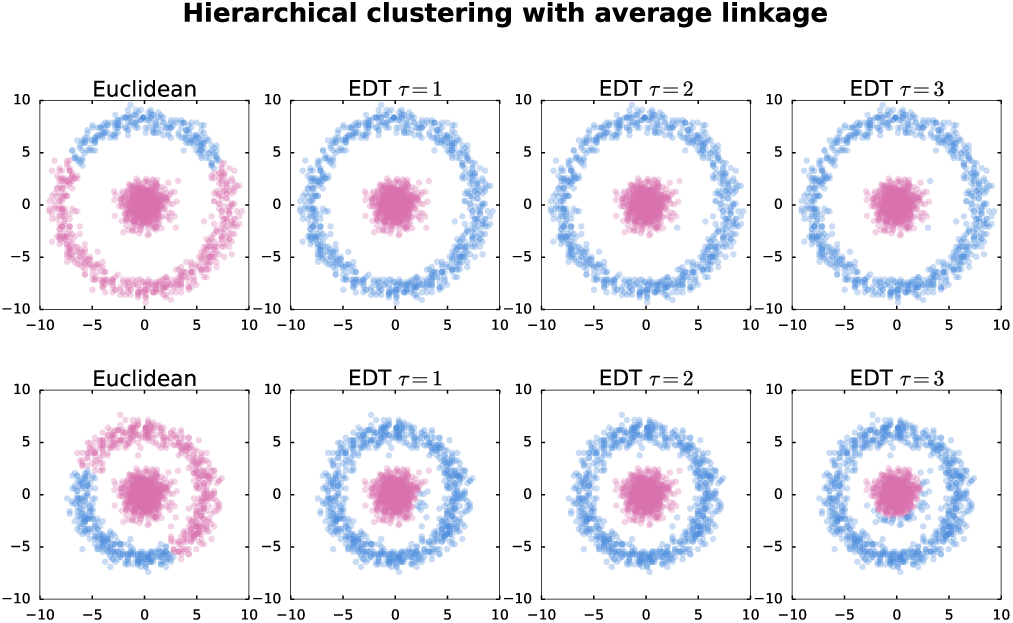
Hierarchical clustering results on “easy” (top) and “hard” (bottom) annulus data sets using Euclidean metric (*τ* = 0) or EDT-enhanced dissimilarities up to three iterations (*τ* = 1, 2, 3) and using average linkage. Dramatic improvements were seen after just one iteration of EDT.

**Figure 9.**
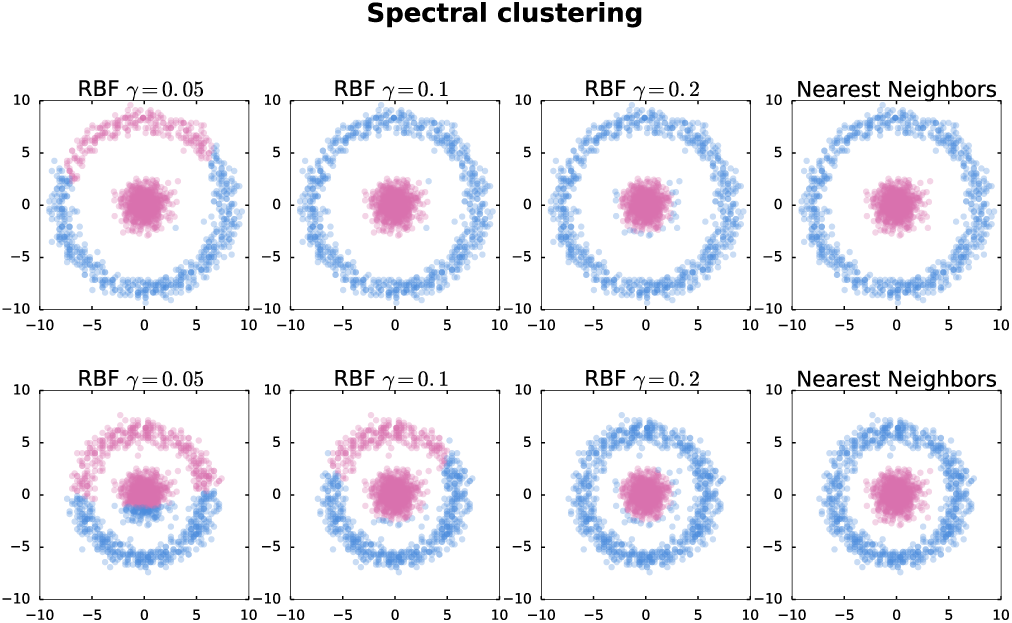
Spectral clustering results for the “easy” (top) and“hard” (bottom) data sets from Fig.8 using the Gaussian RBF kernel exp 
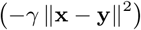
 with γ = 0.05; 0.1 or 0.2 (first 3 columns) and the “nearest neighbors” method (last column) retaining only 10 nearest neighbors of each point to assemble a sparse similarity matrix. In SVM, nonlinear kernels with tunable hyperparameters are usually more powerful than a vanilla linear kernel x · y; however, an unthresholded continuous measure of relatedness between sample points is not necessarily a bless for unsupervised learning algorithms.

**Observation 4** *EDT is able to globally warp the data space on the length scale comparable to inter-cluster distances, such that points far from the majority distribution become approximately identi ed. EDT thus topologically changes*ℝ^*n*^ to *S^n^*.

In application, the EDT will asymptotically group outliers that are very dissimilar to all clusters and may be dissimilar among themselves into one “unclassifiable” cluster in an automatic fashion.

Lastly, we considered the effect of EDT in a probabilistic sense. The initial dissimilarity *d*^(0)^ can be thought of as a random matrix calculated from data sampled from a probability distribution. We replaced the ideal clusters in ℝ^2^ in Fig.1(c) by two independent bivariate Gaussian distributions 
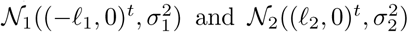
 and 
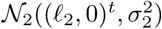
 located symmetrically about the origin, i.e. initially *ℓ*_1_ = *ℓ*_2_. We then placed a test point at the origin and two anchor centroids at *x* = *ℓ*_1_ and *x* = *ℓ*_2_. Denoting the transformed value of *ℓ*_*i*_ after one application of EDT by *ℓ*'_*i*_, we used Monte Carlo simulations to compute the probability 
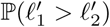
, which may be viewed as the probability that the test point is clustered with 
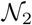
. We performed the calculation in two different settings: (1) 
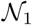
 and 
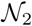
 have same number of samples (*m*_1_ = *m*_2_), but different variances; and (2) 
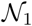
 and 
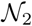
 share the same variance 
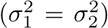
, but different number of samples. We found that the test point was more likely to join (1) a cluster drawn from the distribution with larger variance, consistent with the local deformation effect that absorbs an outlier near the boundary of a cluster into the cluster, or (2) a cluster with fewer samples, consistent with the global deformation effect of EDT that makes points from the majority distribution similar to each other. More precisely, we empirically found the 
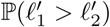
 to be a hyperbolic tangent sigmoid function in *m*_1_/(*m*_1_ + *m*_2_) and −log_2_(σ_1_ / σ_2_), as shown in Fig.2(e-f).

### C. Application of EDT in two gene expression data sets

We tested the power of EDT on two publicly available gene expression data sets: (1) 59 cancer cell lines from NCI60 in 9 cancer types, (2) 116 blood cell samples in 4 cell types from human hematopoietic stem cell differentiation data set [17], with 4,000 most variable genes in each data set as features. We performed hierarchical clustering using the first few iterations of EDT dissimilarity. We used the variation of information (VI) as a well-defined distance between two clustering results [7]; using the given cell types as the reference clustering, we optimized the threshold for cutting the dendrogram into clusters and quantified the performance of clustering with the minimum distance to reference clustering (Fig. 10).

**Figure 10.**
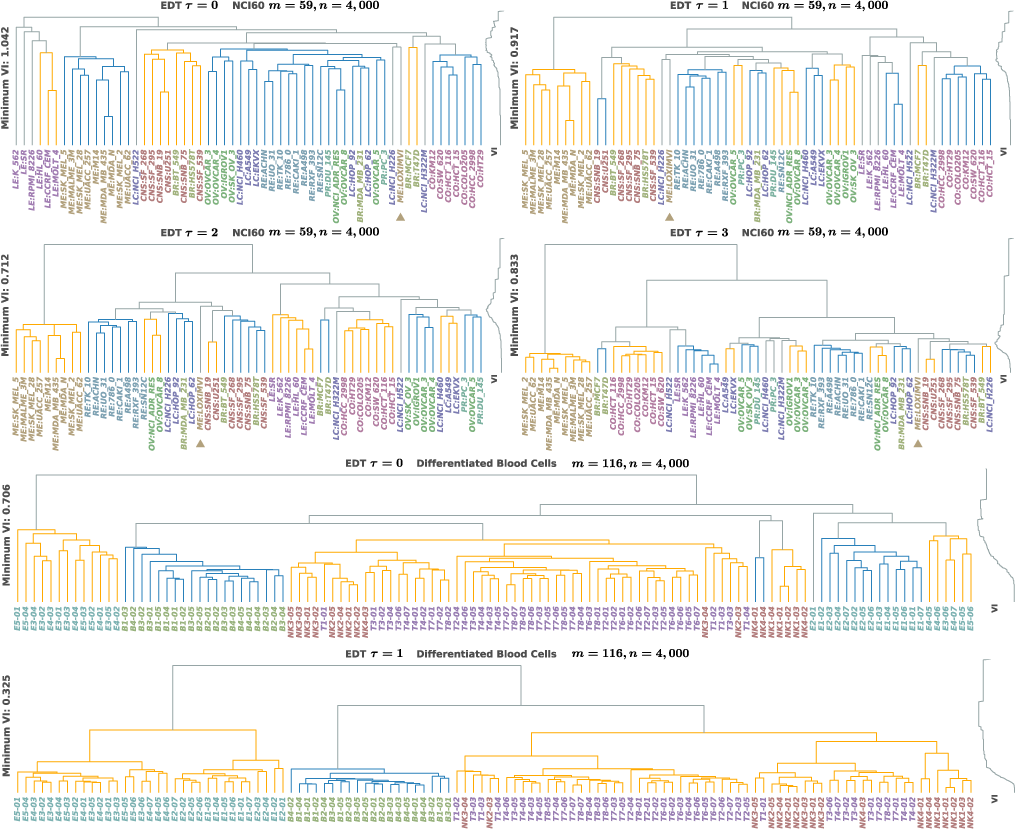
Hierarchical clustering of NCI60 cancer cell lines (row 1-2, *m* = 59 samples) and human differentiated blood cells (row 3-4, *m* = 116 samples) with n = 4,000 most variable genes (with largest standard deviations across all samples) in each each data set. For the NCI60 data, the original Euclidean distance (*τ* = 0) gave minimum VI of 1:042; but, after two rounds of EDT (*τ* = 2), the VI reduced by 31:7% to 0:712. The original Euclidean distance failed to combine all leukemia (LE) cell lines, but EDT (*τ* = 2; 3) brought LE cell lines together into a single cluster. From the very beginning (*τ* = 0), the melanoma cell lines were in a distinct single cluster except for one outlier LOXIM-VI, which is a desmoplastic melanoma cell line and is biologically similar to neuro broma. Among the misclassified cell lines after two iterations of EDT, the LOXIM-VI found itself more similar to the mixture cluster of central nervous system (CNS) and breast cancer (BR) cell lines. For the blood cell data, the original Euclidean distance split the erythrocyte (E*k*, where larger values of *k* indicate latter stages of of maturity) samples into several small sub-clusters, and the VI was 0.706. After one iteration of EDT, the VI reduced by 54.0% to 0.325, and all Ek samples were grouped into a single cluster with two branches – immature red blood cells (E1, E2) and more mature blood cells (E3, E4, E5) – well separated from the immune cells: T-cells, B-cells, and natural killer (NK) cells. These results support that the EDT can help improve clustering performance in real data analysis.

For the NCI60 data, the original Euclidean distance (*τ*= 0) gave minimum VI of 1.042; but, after two rounds of EDT (*τ* = 2), the VI reduced by 31:7% to 0:712 (top two rows in Fig. 10). The original Euclidean distance failed to combine all leukemia (LE) cell lines, but EDT (*τ* = 2, 3) brought LE cell lines together into a single Cluster. From the very beginning (*τ* = 0), the melanoma cell lines were in a distinct single cluster except for one outlier LOXIM-VI. Among the misclassi ed cell lines after two iterations of EDT, the LOXIM-VI found itself more similar to the mixture cluster of central nervous system (CNS) and breast cancer (BR) cell lines; the result is consistent with the fact that LOXIM-VI is a desmoplastic melanoma cell line and is biologically similar to neurofibroma [18].

For the blood cell data, the original Euclidean distance split the erythrocyte (E*k*, where larger values of *k* indicate latter stages of of maturity) samples into several small sub-clusters, and the VI was 0.706 (bottom two rows in Fig. 10). After one iteration of EDT, the VI reduced by 54.0% to 0.325, and all Ek samples were grouped into a single cluster with two branches – immature red blood cells (E1, E2) and more mature blood cells (E3, E4, E5) – well separated from the immune cells (T-cells, B-cells, and, natural killer cells). These results support that the EDT can help improve clustering performance in real data analysis.

## III. DISCUSSION

In this paper, we have developed the notion of effective dissimilarity transformation to enhance the performance of hierarchical clustering, utilizing only the geometric information of all pairwise dissimilarities. The nonlinear transformation adjusts the dissimilarities according to the global distribution of data points. The EDT can be interpreted either as deformation of the feature space or as the result of emergent interactions among all sample points. Specifically, we devised a probe to detect local “tension,” or the force field due to ambient sample points, in a deformed feature space. On a global scale, the EDT is able to change the topology of original Euclidean feature space into a compact sphere. Furthermore, iterating the EDT produces a discrete-time dynamical process purely driven by data set geometry. Using carefully designed Gedankenexperimente, we have shown that EDT has the following properties: (1) perspective contraction, (2) cluster condensation, (3) local deformation, and (4) global deformation effects. These properties arise as different facets of the same mathematical transformation and, thus, should be interpreted in a unified manner. The cosine similarity of EDT is akin to distance correlation [14] and measures the similarity of two random vectors obtained from pairwise similarities to all sample points. Properties (1), (2) and (4) can be understood as mutually enhancing the similarity among a subset of points that share common dissimilar points, while property (3) suggests that common similar points can enhance the similarity between “local” or slightly less similar points.

An adjustable regularizer, such as the number of nearest of neighbors in spectral clustering, is able to qualitatively improve an unsupervised learning algorithm. We have shown that spectral clustering [5] using Gaussian RBF kernels may lead to suboptimal clustering even for some easy synthetic data sets. The reason lies in the fact that Gaussian RBF kernels produce a fully connected network: after restricting each node to communicate with only a specified number of nearest neighbors, the resulting similarity network became sparse and the performance of spectral clustering improved. The sequence of iterated EDT indexed by discrete “time” *τ* plays a similar role in hierarchical clustering: increasing *τ* brings similar sample points into tighter proximity, while enhancing the contrast between clusters (communities). The EDT thus helps hierarchical clustering by utilizing information about the global data distribution. Furthermore, the improvement in clustering accuracy arises from the transformation of data set geometry; thus, any learning algorithm based on pairwise dissimilarity should also benefit from the desirable properties of EDT.

Although the key properties of EDT were first extracted in low feature dimensions in this paper, these advantages, arising from capturing the intrinsic geometry of data distribution, are independent of the feature space dimension, as demonstrated by our finding that EDT also improved the hierarchical clustering of two biological data sets containing 4; 000 features. As an additional verification of the robustness of EDT in high feature dimensions, our simulation shows that the EDT helps increase the contrast in dissimilarity of bimodal Gaussian clouds even in feature dimensions as high as 10^3^, where EDT adapts to the increase in feature dimension by increasing the “time” index *τ* (Appendix B5).

## ACKNOWLEDGMENTS

We thank Alex Finnegan and Hu Jin for critical reading of the manuscript and helpful comments. This research was supported by a Distinguished Scientist Award from Sontag Foundation and the Grainger Engineering Breakthroughs Initiative.

## Appendix A: Data preparation

Two public data sets were used in the hierarchical clustering analysis: (1) NCI60 gene expression data in 59 cancer cell lines comprising 9 cancer types, and (2) 116 differentiated blood cell samples in 4 cell types from human hematopoietic cell (HHC) gene expression data [17]. The 9 cancer types in NCI60 data were 5 breast (BR), 6 central nervous system (CNS), 7 colon (CO), 6 leukemia (LE), 10 melanoma (ME), 8 non-small cell lung (LC), 7 ovarian (OV), 2 prostate (PR), and 8 renal (RE) cancer. The 4 cell types in the HHC data were red blood cells or erythrocytes (E), T-cells (T), B-cells (B), and natural killer cells (NK). For both data sets, approximately four thousand most variable genes were selected as features. Samples from each data set were first clustered with the usual Euclidean distance 
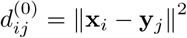
, and then with the EDT dissimilarity 
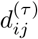
 computed from 
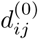
. Average linkage was used in all hierarchical clustering analysis, unless indicated otherwise. To quantify the clustering performance unambiguously, the minimum distance from a given clustering to the standard reference clustering was found by measuring the variation of information (VI), which is a well-defined metric function that computes the distance between different partitions (clusterings) of a given set [7]. The reference clustering for NCI60 was the known 9 cancer types (BR, CNS, CO, LE, ME, LC, OV, PR, and RE); the reference clustering for HHC was the known 4 cell types (E, T, B, and NK).

## Appendix B: Effective dissimilarity transformation

The following properties of EDT mentioned in the main article were obtained from the Gedankenexperimente illustrated in Fig.2(b-f).

### 1. Perspective contraction

The 3 points {*P*_1_, *P*_2_, *P*_3_} shown in Fig.1(b) form two distinct configurations: (1) aligned in a line, and (2) forming a triangle in a plane. For case (1), let *P*_1_ and *P*_2_ be at *x* = +*a*/2 and −*a*/2, respectively, and *P*_3_ at *x* = *b* > *a*/2. Then, the original dissimilarity matrix is

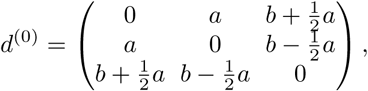

and the transformed feature vectors are:

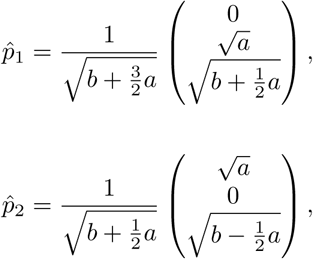

and

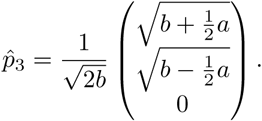

From these feature vectors, we compute the first EDT dissimilarity matrix components to be

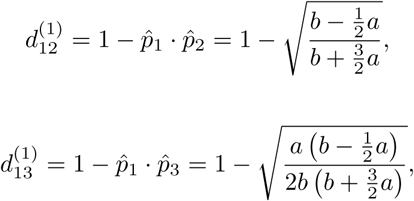

and

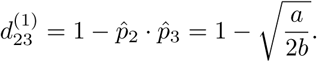

As 
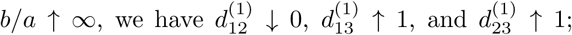
 in other words, the EDT ruler length will shrink to zero if the observer moves away from the ruler. Next, we can calculate the relative dissimilarity, i.e. the observed difference between *P*_1_ and *P*_2_ from the perspective of *P*_3_ measured in units of the transformed dissimilarity 
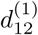
, to be

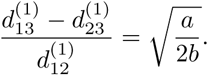

Therefore, as the observer moves away from the ruler, the EDT ruler length shrinks to zero, but the observed difference shrinks even faster. In the application of hierarchical clustering, the diminishing difference between the nearest (*P*_2_) and the farthest (*P*_1_) point with respect to the outlier *P*_3_ implies that clustering derived from the EDT tends to be robust against the choice of linkage, which may be single (nearest point), average, or complete (farthest point).

For case (2), we set up a Cartesian coordinate system in ℝ^2^such that *P*_1_, *P*_2_, and *P*_3_ are located at (0, *a*/2); (0,−*a*/2), and (*b*, 0), respectively, where we assume *a, b* > 0: The original Euclidean distance matrix is thus

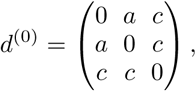

Where 
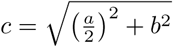
. The transformed feature vectors are

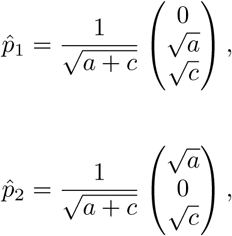

and

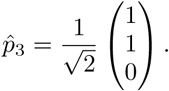

The corresponding transformed dissimilarity matrix elements are

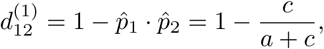

and

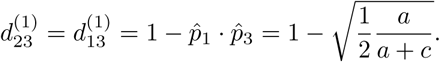

As *b/a* increases to infinity, 
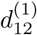
 monotonically decreases to zero. Thus, the effective ruler length *d*_12_ approaches 0 from the perspective of point *P*_3_ as it moves far away.

### 2. Cluster condensation

When clustering real datasets, the contrast between the inter-cluster distance and the intra-cluster variance is often not very dramatic, making it very diffcult to separate the clusters. Therefore, if the data points could condense to the respective centroid locations, then it would improve clustering accuracy considerably; this effect is precisely what EDT accomplishes. For the synthetic data shown in Fig.1(c), the EDT centroid-centroid dissimilarity 
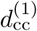
 increased relative to the average centroid-satellite distance 
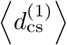
, or the contrast captured by the ratio 
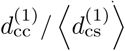
, grew more rapidly than 
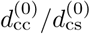
. Moreover, for fixed 
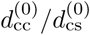
, increasing the number of satellites *m_s_* around each centroid amplified the contrast ratio (Fig. 2(a)).

Throughout the simulations, we did not use any information about the cluster labels, and the improvement of contrast is purely driven by the data. The dense clusters condense while pushing themselves away from other clusters. In other words, within a cluster, the EDT acts similar to gravity, whereas the transformation inflates the space between clusters.

### 3. Local deformation

In Fig.1(e) *r* denotes the radius of the cluster and *ℓ* the distance between a single test point and the cluster centroid. We simulated the effect of increasing *ℓ* on the EDT. As 
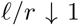
, we observed a window where the transformed ratio 
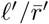
 was less than or equal to the local peak at *ℓ*/*r* = 1 (Fig.2(b)). This phenomenon can be interpreted as the cluster’s trying to reabsorb the test point that is escaping to become an outlier. We also observed that the range of absorption window increased as the cluster size *m_s_* increased, thus making it easier for an outlier to tunnel back to a denser cluster (Fig.2(b)). Moreover, the test point also deformed the shape of the cluster, and the satellite points on the circle acted like an elastic membrane that trapped the test point and hindered it from escaping the cluster through elongation.

### 4 Global deformation

Consistent with the single test point example, the cluster in Fig.1(f) tended to attract the two escaping outliers, as manifested by the fact that as *m_s_* increased, the ratio *ℓ*'/*r*' decreased (Fig.2(c)). Counterintuitively, 
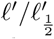
 also dropped below 1 as *ℓ*/*r* increased (Fig.2(d)); that is, the two test points became more similar as they departed from the cluster centroid in opposite directions. This paradox can be resolved by merging the points at infinity to a single point, or by topologically transforming the Euclidean space into a hypersphere. We explicitly demonstrated our idea using two circularly distributed data sets shown in Fig. 7. We first observed that the effective dissimilarity between two neighboring points in the outer rings shrank faster than that between neighboring points in the inner rings. To better visualize this phenomenon, we then displayed the dissimilarities on a sphere using the following methods.

For a ring of *k* points distributed on a unit 2-sphere at constant colatitude *θ* ∈ [0,π] and uniformly partitioned longitude *ϕ*_*i*_ ∈ [0,π] *i* = 1,…, *k*, the latitude distance *ℓ* between any two neighboring points is equal to sin *θ δϕ*, where *δϕ* = 2π/*k*. Thus, *ℓ* attains its maximum value *ℓ*_max_ = *δϕ* at the equator 
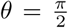
. Note that regardless of the size of *δϕ*, we always have *ℓ*=*ℓ*_max_ = sin *θ* we will utilize this fact to display the EDT-deformed concentric rings shown in Fig. 7. For this purpose, it might appear natural to identify the centroid as the north pole of the sphere, and then identify the colatitude *θ*' of a ring as the EDT dissimilarity between the centroid and a point on the ring. However, while the distance between two neighboring data points on the sphere at such *θ*' would then be fixed to be sin *θ*' *δϕ*, the actual EDT dissimilarity *ℓ*' might be different. We thus empirically calculated the function *f*(*θ*') that satisfies *ℓ*'=*f*(*θ*')*ℓ*'_max_. We then used the location 
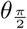
 of the global maximum of *f* to calibrate the equator location, and then calculated the effective colatitudede 
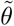
 defined as 

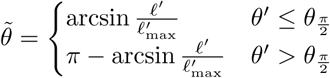

to display the concentric rings on the sphere, as shown in Fig.7. Fig.11 shows the *f*(*θ*') for the two circular data sets shown in Fig. 7 after *τ* iterations of EDT.

**Figure 11.**
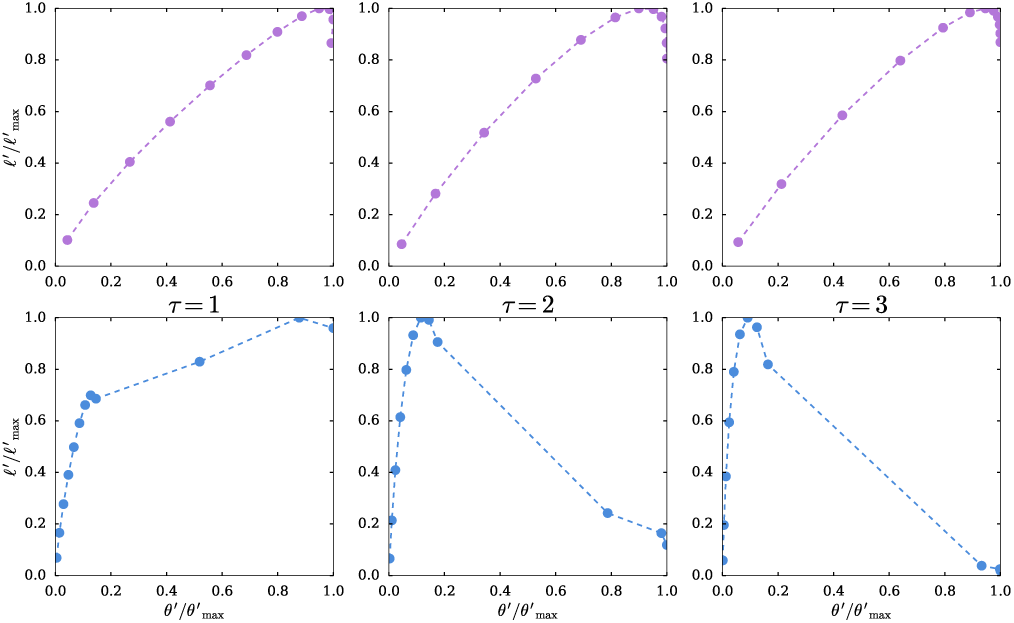
Plots of the empirical function *f* that satisfies 
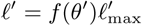
, where *ℓ*' is the EDT dissimilarity between two neighboring points on a circle and *θ*' is the EDT dissimilarity between the centroid and the circle. The three plots on the top (bottom) correspond to the top (bottom) three spheres in Fig.7

### 5 EDT and the curse of dimensionality

The loss of contrast in Euclidean distance is one of the symptoms of the curse of dimensionality; to be exact, the longest distance 
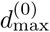
 and shortest distance 
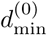
between any pair of points in a data set will both asymptotically approach the mean distance 
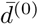
 in the large feature dimension limit 
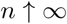
 To see whether EDT can help improve the contrast between clusters in high dimensions, we simulated two *n*-dimensional Gaussian distributions 
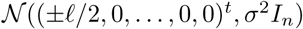
100 points from each, for *ℓ*σ= 0; 4; and 10. We then computed the Euclidean distance matrix *d*^(0)^ and subsequent effective dissimilarity matrices 
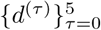
, Fig. 12 shows the normalized pairwise maximum 
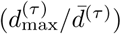
 and minimum 
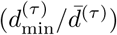
 distance between data points in each dimension. The difference 
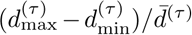
 generally became larger as the EDT index *τ* increased, and the improvement in contrast over the original Euclidean distance in high dimensions was very dramatic when *ℓ*≫σ, as seen for *n* = 1000 in Fig. 12(c).

**Figure 12.**
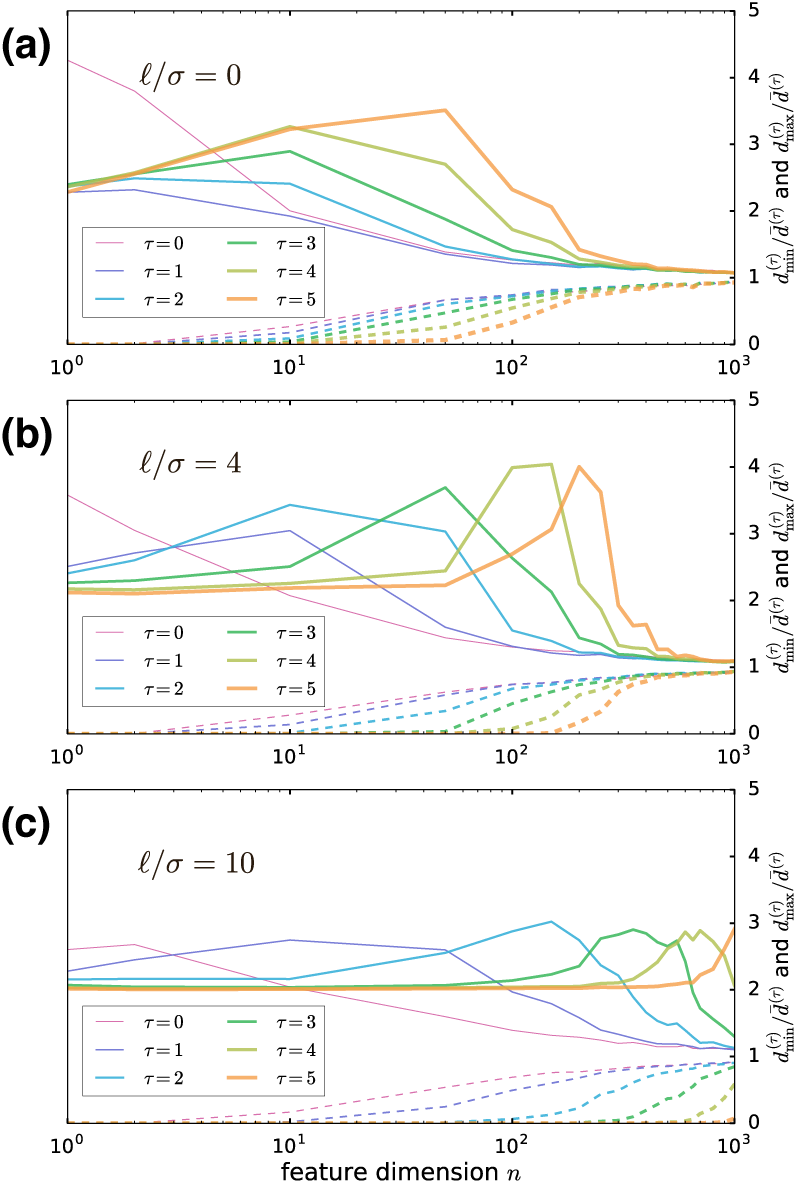
Plots of maximum and minimum dissimilarities normalized by mean dissimilarity: 
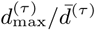
 (solid) 
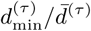
 (dashed) of two multivariate normal distributions 
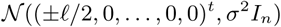
 in ℝ^*n*^with variations in (a) *ℓ*σ = 0, (b) *ℓ*σ/ = 4, and (c) *ℓ*σ/ = 10. For all three cases (a-c), EDT (*τ*>0) enlarged the difference between 
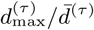
 and 
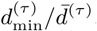
, and hence enhanced the contrast; when the initial inter-cluster distance *ℓ*≫σ, EDT with high index *τ* preserved contrast dramatically relative to initial Euclidean distance *d*^(0)^, consistent with cluster condensation effect of EDT.

